# Evaluating permutation-based inference for partial least squares analysis of neuroimaging data

**DOI:** 10.1101/2024.08.02.606412

**Authors:** Matthew Danyluik, Yashar Zeighami, Alice Mukora, Martin Lepage, Jai Shah, Ridha Joober, Bratislav Misic, Yasser Iturria-Medina, M. Mallar Chakravarty

**Affiliations:** Cerebral Imaging Centre, Douglas Mental Health University Institute, Montreal, Canada; Integrated Program in Neuroscience, McGill University, Montreal, Canada; Department of Psychiatry, McGill University, Montreal, Canada; Prevention and Early Intervention for Psychosis, Douglas Mental Health University Institute, Montreal, Canada; Department of Psychology, McGill University, Montreal, Canada; McConnell Brain Imaging Centre, McGill University, Montreal, Canada; Department of Neurology and Neurosurgery, McGill University, Montreal, Canada; Ludmer Centre for Neuroinformatics and Mental Health, McGill University, Montreal, Canada; Department of Biological and Biomedical Engineering, McGill University, Montreal, Canada

**Keywords:** Partial least squares, permutation testing, statistical inference, multivariate analysis

## Abstract

Partial least squares (PLS) is actively leveraged in neuroimaging work, typically to map latent variables (LVs) representing brain-behaviour associations. LVs are considered statistically significant if they tend to capture more covariance than LVs derived from permuted data, with a Procrustes rotation applied to map each set of permuted LVs to the space defined by the originals, creating an “apples to apples” comparison. Yet, it has not been established whether applying the rotation makes the permutation test more sensitive to whether true LVs are present in a dataset, and it is unclear if significance alone is sufficient to fully characterize a PLS decomposition, given that complementary metrics like strength and split-half stability may offer non-redundant information about the LVs. Accordingly, we performed PLS analyses across a range of simulated datasets with known latent effects, observing that the Procrustes rotation systematically weakened the null distributions for the first LV. By extension, the first LV was nearly always significant, regardless of whether the effect was weak, undersampled, noisy, or simulated at all. But, if no rotation was applied, all possible LVs tended to be significant as we increased the sample size of UK Biobank datasets. Meanwhile, LV strength and stability metrics accurately tracked our confidence that effects were present in simulated data, and allowed for a more nuanced assessment of which LVs may be relevant in the UK Biobank. We end by presenting a list of considerations for researchers implementing PLS permutation testing, and by discussing promising alternative tests which may alleviate the concerns raised by our findings.

## 1. Background

The partial least squares (PLS) correlation is a popular technique in neuroimaging research, typically used to identify sets of brain and behavioural measures that covary across a sample, referred to as latent variables (LVs) (Abdi et al., 2013; Krishnan et al., 2011; McIntosh & Lobaugh, 2004; McIntosh & Mišić, 2013). PLS was introduced to the neuroimaging community as a means of finding the overlapping information across a *pattern* of brain measures and linking it to outcome variables of interest – a multivariate extension of mass univariate approaches (McIntosh et al., 1996). In turn, PLS both capitalizes on the “many to many” functional organization of the brain and is theoretically well-suited for finding robust brain mappings in the high-dimensional, multimodal datasets growing in popularity across the literature (Genon et al., 2022; Mihalik et al., 2022).

In a PLS analysis, LVs are considered statistically significant if pass permutation testing – in other words, if they tend to explain more covariance than LVs derived from randomly shuffled data (McIntosh & Lobaugh, 2004). Importantly, during this procedure, a Procrustes rotation is applied to each set of permuted LVs to align them as closely as possible with those from the original decomposition (McIntosh & Lobaugh, 2004; Milan & Whittaker, 1995). As such, permutation testing can be thought of as comparing each latent variable with permuted latent variables capturing similar brain/behaviour associations (for more details, see *Methods*).

While the Procrustes rotation is a standard component of a neuroimaging PLS workflow, the impact of this design choice on LV significance relative to the unrotated condition has not been systematically explored in the PLS literature. Moreover, the rotation is usually only applied to the smaller/behavioural component of the LVs, and it is unclear if this achieves similar results as rotating the larger/brain component, or rotating both, since PLS is fundamentally a two-sided procedure. As initial observations from our group suggested that different permutation tests (unrotated, or rotating brain/behaviour/both) performed on the same dataset can give radically different *p*-values (see *Supplementary Materials, Section S2* for an overview), in the present study, we sought to determine whether each rotation method could reliably detect ground truth effects in simulated data, and whether they would lead to different conclusions regarding LV significance in samples drawn from the UK Biobank.

Finally, while LVs may be significant by permutation testing, it is well established that effect size estimates are crucial for making neurobiological inferences about a finding (Chen et al., 2017), and there are growing concerns that the results from PLS and other multivariate techniques do not necessarily reproduce in held-out samples (Churchill et al., 2013; Nakua et al., 2024; Ji et al., 2021; Helmer et al., 2024; Dinga et al., 2019), regardless of significance (Kovacevic et al., 2013; McIntosh, 2022). Accordingly, we also evaluated whether alternative PLS outcome metrics, namely LV strength (covariance explained) and stability (across sample splits), offered information beyond significance about LV quality in both simulated and UK Biobank data. Together, then, we aimed to establish whether significance alone is sufficient to fully characterize a PLS decomposition.

## 2. Methods

### 2.1. The partial least squares correlation

First, we introduce how PLS is performed in the case of implementing a PLS correlation to analyze brain-behaviour covariance, sometimes referred to as behavioural PLS (Krishnan et al., 2011). Assume we have brain matrix *X* and behavioural matrix *Y*, each with *n* rows of observations and with z-scored feature vectors in the columns. To implement PLS, the cross-correlation matrix between *X* and *Y* is subjected to a singular value decomposition (SVD):

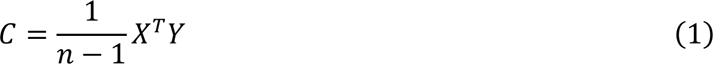

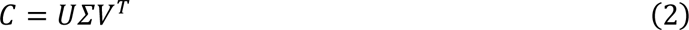

With the left and right singular vectors of *C* in the columns of *U* and *V*, respectively, and the singular values across the diagonal of *∑*. A given latent variable consists of matching columns of *U* and *V* (interpreted as brain and behavioural feature weights, respectively) and their corresponding singular value (proportional to covariance explained). Latent variables are mutually orthogonal and are ranked by their singular value, and the number of latent variables corresponds to the rank of *C* (typically, the lesser of the number of columns in *X* or *Y*). For more details, see McIntosh et al. (1996).

### 2.2. Permutation testing and the Procrustes rotation

Next, we describe how PLS permutation testing is normally implemented, with a Procrustes rotation applied to the smaller (“behavioural”) component of the decomposition. First, PLS is performed as described above:

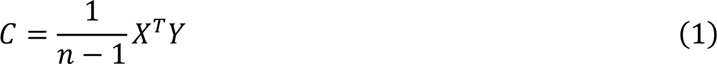

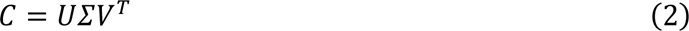

Then, if we consider a single permutation instance where the rows of *Y* are permuted, giving **Y**^(*p*)^, and PLS is repeated:

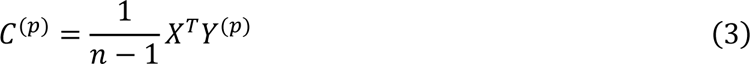

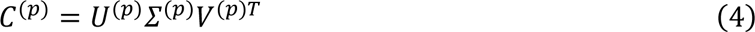

The orthogonal Procrustes problem (Schönemann, 1966) is solved to find the orthonormal matrix *R* which, when applied to *V*^(p)^, maps it as closely as possible to *V* in a least squares sense (McIntosh & Lobaugh, 2004):

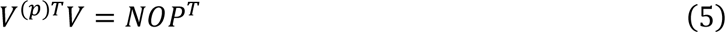

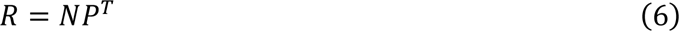

Where *N*, *O*, and *P* are the result of an SVD. For a full proof and discussion, see Schönemann (1966). Next, each row of *R* is scaled by its corresponding diagonal element of *∑*^(p)^, and the transformation is applied to rotate (and/or reflect) the columns of *V*^(p)^:

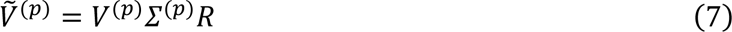

The transformation can both reorder the singular vectors and redistribute covariance explained across them (McIntosh & Lobaugh, 2004). Then, the L2 norms of the scaled and transformed singular vectors are calculated:

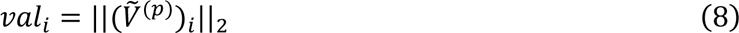

With *i* representing any given column of 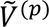. The values are added to the null distributions for their corresponding latent variables (the first value for LV1, the second for LV2, etc.), and the procedure is repeated on each permutation instance. Finally, the singular values from the original decomposition, across the diagonal of *∑*, are tested against their respective null distributions, giving *p*-values for each corresponding to the proportion of permuted singular values exceeding the original. Typically, LVs with *p* < 0.05 are considered statistically significant.

### 2.3. Simulated data

The following analyses are summarized in *Fig. 1*.

**Figure 1:**
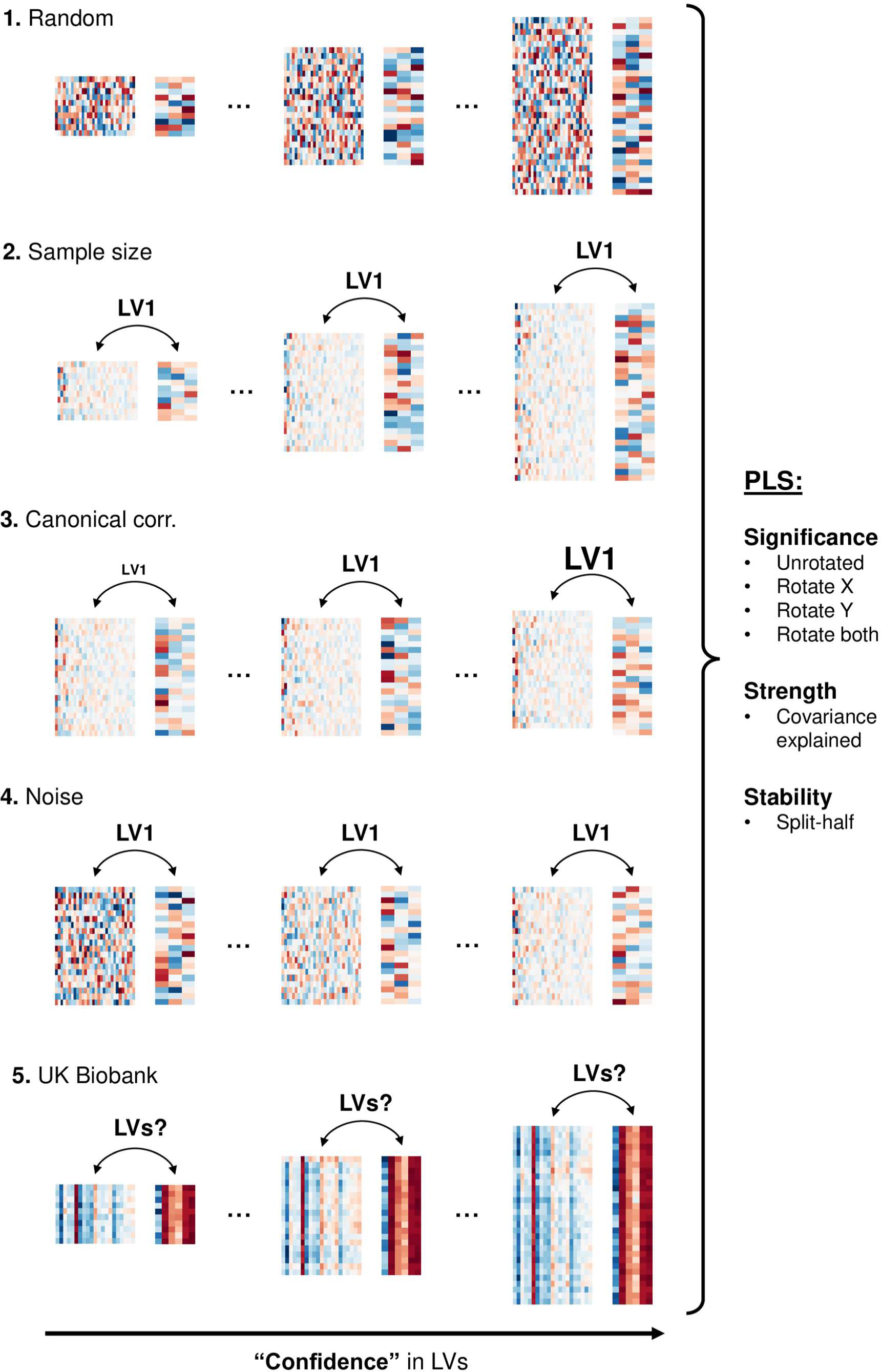
Workflow for the simulated and UK Biobank analyses. First **(1)**, we generated datasets of values randomly drawn from a normal distribution, expecting our PLS outcome metrics to indicate that no effects were present (i.e., that LV1 from these datasets would be insignificant, weak, and unstable). Next, we generated a series of datasets with increasing “confidence” that a simulated LV1 was present, corresponding to **(2)** a larger sample, **(3)** a stronger effect, or **(4)** less noise. After performing PLS on each dataset, we expected that the listed outcome metrics would track our confidence, such that LV1 from the rightmost datasets would be stronger, more significant, and more stable than LV1 from the leftmost datasets. Finally **(5)**, we varied the size of subsamples drawn from our broader pool of UK Biobank participants, offering progressively more evidence for the “true” effects that may have been present. After performing PLS on each dataset, we evaluated whether significance, strength, and stability depended on sample size for various LVs of interest.

#### 2.3.1. Randomly generated data

First, we aimed to evaluate whether PLS could “find” latent effects in randomly generated datasets without any explicitly coded associations between the *X* and *Y* matrices. We created 100 different datasets at each of 10 sample sizes, with sample sizes log-spaced between 10 and 10 000. All samples were drawn from a normal distribution with mean 0 and variance 1. The number of features in both matrices was held constant across datasets (90 *X*, 10 *Y*).

After performing PLS on each dataset, we calculated the LV1 pass rate at each sample size, corresponding to the proportion of the 100 datasets with *p* < 0.05 for LV1. In this case, the pass rate was our proxy for the permutation test error rate, and we directly compared pass rates between rotation methods (see *PLS implementation* below more details). LV1 strength and stability were also calculated at each sample size, allowing us to assess which PLS outcome metrics best reflected the ground truth of no true between-set associations (see also *PLS implementation* below).

#### 2.3.2. Data with ground truth effects

Next, we tested the sensitivity of LV significance, strength, and stability to the signal-to-noise ratio of known simulated effects. We generated a series of datasets, each with one true LV present. However, our “confidence” in the effect varied across datasets, with more confidence corresponding to a larger sample, a stronger effect, and less noise. After performing PLS on each dataset, we could evaluate which outcome metrics best tracked our confidence in LV1, and by extension were most sensitive to the dataset’s key properties (see *Fig. 1* for a summary).

To create our *X* and *Y* matrices, we used the generative modeling of multivariate relationships (GEMMR) Python package, version 0.2.6 (Helmer et al., 2024). GEMMR allows users to specify the properties of a correlation matrix *C*, including the correlation matrices *C*_XX_ and *C*_YY_ and the cross-correlation matrix *C*_XY_, then generate a dataset by drawing a defined number of samples from a multivariate normal distribution with mean 0 and correlation matrix *C*.

Specifically, since PLS is the SVD of a cross-correlation matrix (see *Eq. (1)* and *(2)*), GEMMR defines *C*_XY_ by generating semi-random *X* and *Y* singular vectors and user-defined singular values (expressed as a canonical correlation) quantifying the ground truth relationships between singular vector pairs. Following Helmer et al. (2024), in our simulations, we only defined the canonical correlation for the first singular vector pair, meaning that further pairs were not constrained to covary with each other. For more details on the simulation protocol and its assumptions, see Helmer et al. (2024).

In the three analyses described below, we generated datasets using GEMMR and tuned a different parameter across simulations (either the sample size, canonical correlation, or noise). All unmentioned parameters were set to their default values.

### Sample size

As above, we generated 100 different datasets at each of 10 sample sizes (i.e., the number of draws from the multivariate normal distribution defined by GEMMR), with sample sizes log-spaced between 10 and 10 000. All other dataset properties were identical, including the number of latent effects (*m* = 1), the canonical correlation of the effect (*r* = 0.3), and the number of features in either matrix (90 *X*, 10 *Y*).

Again, after running PLS on each dataset, we calculated the LV1 pass rate at each sample size, quantifying whether a permutation test could consistently detect a known simulated effect. LV1 pass rates were calculated for each permutation test method and contrasted with both strength and stability.

### Canonical correlation

Next, for each of 10 canonical correlations (proportional to ground truth singular values of the first latent effect), we generated 100 different datasets, with the canonical correlations linearly spaced between 0.1 and 0.9. The number of latent effects (*m* = 1), number of features (90 *X*, 10 *Y*), and sample size (*N* = 1000) were held constant across datasets. LV1 pass rates, alongside strength and stability metrics, were calculated in turn.

### Noise

Finally, for *X* and *Y* matrices generated by GEMMR, we added equally sized matrices of Gaussian noise after simulation. Specifically, we generated 100 different datasets at each of 10 noise levels, with the noise level defined by the standard deviation of a Gaussian (the 10 standard deviations were linearly spaced between 0 and 1). As above, we held all other parameters constant across datasets, including the number of latent effects (*m* = 1), the canonical correlation of the effect (*r* = 0.3), number of features (90 *X*, 10 *Y*), and sample size (*N* = 1000). LV1 pass rates (by rotation method), strength, and stability were compared accordingly.

#### 2.3.3. PLS implementation

For a given dataset (a pair of simulated *X* and *Y* matrices), PLS was performed using the *behavioral_pls* function from the pyls library (https://github.com/rmarkello/pyls/) in Python 3.7, which mirrors the original MATLAB implementation for neuroimaging (https://www.rotman-baycrest.on.ca/index.php?section=84). We performed 4 permutation tests per dataset, each with different rotation methods, described below. Note that the “brain” and “behaviour” designation is arbitrarily based on relative matrix size.

- *None*: the rows of *Y* were permuted and no rotation was applied to the permuted latent variables. Their singular values were directly added to the null distributions.
- *Behaviour*: as a proxy for the typically smaller behavioural matrix, the rows of *Y* were permuted, and the Procrustes rotation was applied to its corresponding singular vectors to generate values for the null distributions.
- *Brain*: as a proxy for the typically larger brain matrix, the rows of *X* were permuted, and the Procrustes rotation was applied to its corresponding singular vectors to generate values for the null distributions.
- *Both*: the rows of *Y* were permuted and the Procrustes rotation was applied to both sets of singular vectors. The values returned from the two procedures were averaged together, then added to the null distributions.

10 000 permutations were performed for each permutation test, and the indices used to shuffle the matrices on a given permutation instance were identical across tests. As such, any differences in the null distributions, and by extension the *p*-values, between the permutation tests were driven by the rotations themselves rather than the reshuffling procedure.

Finally, to complement significance testing, we assessed both the covariance explained and split-half stability of the LVs. Regarding split-half stability, we employed the metric proposed by McIntosh (2022) and Nakua et al. (2024). Specifically, for a given dataset, we randomly split the sample into two halves and ran PLS on each half separately. Then, for each LV, we calculated the Pearson correlation between the singular vectors from the two halves separately for “brain” (*U*) and “behaviour” (*V*). We took the absolute value of each correlation coefficient to account for arbitrary sign flips and repeated the procedure 100 times.

### 2.4. UK Biobank

Next, we tested whether the *p*-values from the different rotation methods would disagree with each other, or diverge from strength and stability metrics, in real neuroimaging data.

Accordingly, we gathered brain and behavioural data from 28 804 UK Biobank participants, with the brain matrix *X* composed of cortical thickness in 64 regions and the behavioural matrix *Y* composed of 17 lifestyle risk factors linked to adverse aging (see *Supplementary Materials, Section S4* for a full description of the dataset).

Then, we drew 100 different subsamples at each of 10 sample sizes from the broader pool of 28 804 participants, with the sample sizes log-spaced between 50 and 20 000 (the low bound ensured that split-half samples were sufficiently variable, and the high bound leveraged most of the participants while still allowing for differences between samples). After performing PLS on each subsample, we evaluated whether significance (for each rotation method), strength, and stability depended on sample size for different LVs, as described in the previous section. The workflow is visualized in the bottom panel of *Fig. 1*. Finally, informed by our full set of simulated and UK Biobank results, we performed a PLS analysis on all 28 804 participants, and explored which measures could conceivably be used as “stopping” metrics for identifying potentially meaningful LVs.

#### 2.4.1. PLS implementation

For a given subsample, PLS was implemented as described in *Section 2.3.5*, with *p*-values calculated for the 4 rotation methods across 10 000 permutations. As above, covariance explained and split-half stability were also assessed for each decomposition.

Finally, for the full dataset of 28 804 participants, we implemented bootstrap resampling to assess LV feature weight reliability. Specifically, participants were sampled with replacement 10 000 times and PLS was performed on each bootstrapped sample. For each LV, we assessed bootstrap ratios for the brain features, corresponding to the ratio between a feature’s mean weight and standard deviation across samples (analogous to a z-score). For the behavioural variables, we assessed loadings, or the correlation between the variable and latent variable scores across participants (with 95% confidence intervals calculated across bootstrap samples). Note that a Procrustes rotation is also typically applied to the brain component of the latent variables during bootstrap resampling (McIntosh & Lobaugh, 2004). We kept this convention for our analysis and did not assess the impact of different rotation methods on bootstrapped feature weights.

## 3. Results

### 3.1. Simulated data

First, we aimed to establish whether the *p*-values from rotated and unrotated PLS permutation tests accurately reflect the presence of known latent effects in simulated data, and whether complementary outcome metrics, like latent variable strength and stability, offer information beyond significance regarding a dataset’s ground truth. Accordingly, we performed PLS on a range of randomly generated datasets, assessing which measures (significance, strength, and stability) correctly indicated that no effects were encoded. Then, we leveraged the GEMMR Python package (Helmer et al., 2024) to simulate a range of datasets with one true latent variable present. Importantly, our confidence in the effect differed across datasets: either the sample size, effect strength, or amount of noise was allowed to vary, with all other parameters held constant. After performing PLS on each dataset, we could then determine which metrics consistently scaled with the quality of the effect (see *Fig. 1* for a summary).

Note that throughout our simulated data analyses, the larger matrix is arbitrarily referred to as “brain”, while the smaller matrix is arbitrarily referred to as “behaviour”, consistent with the conventions commonly used in neuroimaging PLS analyses.

#### 3.1.1. Analysis 1: Random data

First, we asked whether our permutation tests were prone to erroneously detecting latent effects in randomly generated data. We simulated 100 different datasets at each of 10 sample sizes, with each sample randomly drawn from a normal distribution. Here, we found that rotated permutation tests considered LV1 significant at all sample sizes (except when the rotation was applied to the smaller “behavioural” matrix, which was less prone to errors at smaller samples), while the unrotated tests rarely detected an effect (*Fig. 2A*). Meanwhile, LV1 strength and stability metrics were consistently low regardless of sample size (*Fig. 2A*).

**Figure 2:**
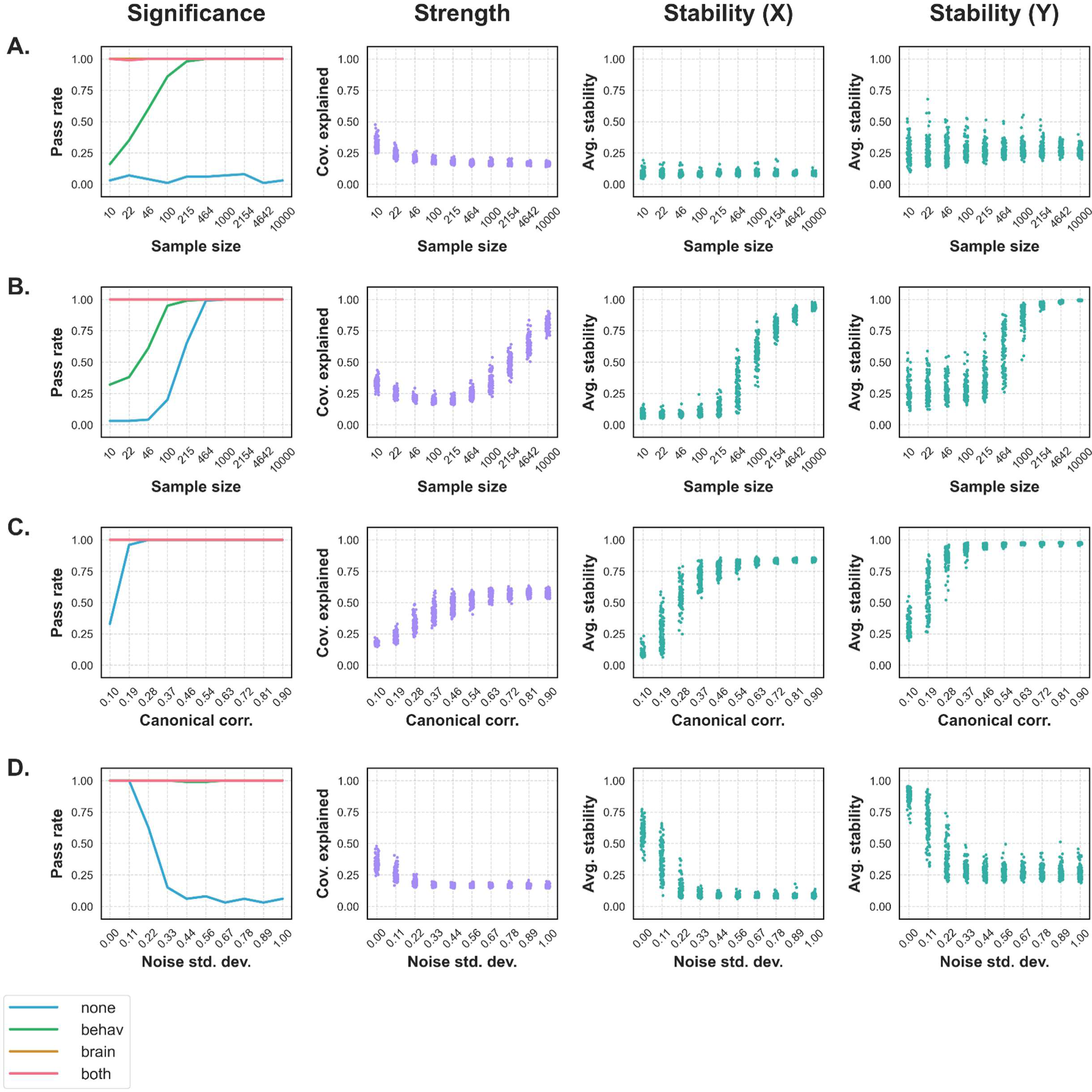
LV1 significance, strength, and stability metrics across each simulated data analysis. The **Significance** column shows the pass rates for LV1, or the proportion of the 100 datasets with a significant (*p* < 0.05) LV1 at each step, plotted separately for each rotation method. The **Strength** column shows the covariance explained of LV1, and the **Stability (X)** and **Stability (Y)** columns show the average split-half stability of LV1 across 100 splits for the “brain” and “behaviour” LV components, respectively. Individual points represent values for unique simulated datasets. **A. (Random data)** Rotated permutation tests consistently deemed LV1 significant, even though no effects were encoded. Meanwhile, the unrotated test was far less likely to find the unsimulated “effect”, and strength and stability metrics remained low regardless of sample size. **B. (Sample size)** Pass rates steadily increased with *N* when no rotation was applied, while all rotated permutation tests were more likely to detect the effect in smaller samples. Like the unrotated pass rates, LV1 strength and stability both climbed with sample size. **C. (Canonical correlation)** All rotated permutation tests consistently detected the first latent variable regardless of how strong it was, while the unrotated test was less sensitive to the weakest effects. Meanwhile, strength and stability both increased with the simulated canonical correlation. **D. (Noise)** The unrotated permutation test failed to consistently detect the simulated effect under noisy conditions, while all rotated tests generally considered LV1 significant regardless of noise levels. Alongside unrotated significance, the strength and stability of LV1 all decreased as noise levels increased.

#### 3.1.2. Analysis 2: Sample size

Next, we asked whether PLS would consider a known latent effect to be more significant, stronger, or more stable in larger samples. As above, we simulated 100 different datasets at each of 10 sample sizes, though each had one latent effect encoded. Here, we observed that all rotated permutation tests were more likely to detect LV1 in relatively small samples than unrotated tests, while the methods converged as *N* increased (*Fig. 2B*). Meanwhile, the covariance explained and split-half stability of LV1 both required a relatively large sample size to stabilize (notably, covariance explained did not increase monotonically with sample size until the number of samples exceeded the number of features) (*Fig. 2B*).

#### 3.1.3. Analysis 3: Canonical correlation

We also tested whether our PLS outcome metrics were sensitive to the strength of a simulated effect. Accordingly, we scaled the canonical correlation of our simulated latent effects, analyzing 100 different datasets at each of 10 canonical correlations. Here, all rotated permutation tests consistently detected LV1 regardless of its strength, while unrotated tests were less likely to detect relatively weak effects (*Fig. 2C*). Predictably, covariance explained climbed with the simulated canonical correlation, and split-half stability did not stabilize until the effect was relatively strong (*Fig. 2C*).

#### 3.1.4. Analysis 4: Noise

Finally, we assessed whether noise would compromise the sensitivity of PLS to simulated effects. Analyzing 100 different datasets at each of 10 noise levels, we observed that all rotated permutation tests consistently detected LV1 regardless of the amount of noise, while the unrotated tests were more conservative in noisier datasets (*Fig. 2D*). Alongside unrotated significance, the covariance explained and split-half stability of LV1 both decreased as noise levels increased (*Fig. 2D*).

#### 3.1.5. Summary

Together, rotated permutation tests were characterized by high error rates in randomly generated data, regardless of which LV component was rotated (“behaviour”, “brain”, or both). Rotated tests also consistently detected LV1, whether the effect was present in a small sample, weak, or obscured by noise, as well as LV2, which was not simulated in any analysis (*Supplementary Fig. S8*). Meanwhile, unrotated *p*-values, covariance explained, and split-half stability accurately scaled with the signal-to-noise ratio of simulated effects and did not suggest that true LVs were present in null data. Further simulated data analyses are shown in *Supplementary Materials – Section S2*.

### 3.2. UK Biobank

Above, we saw that rotated permutation tests tended to systematically pass LV1, regardless of its strength, stability, or simulated properties. Accordingly, we wondered whether similar limitations could be observed in real neuroimaging data. Specifically, we analyzed brain (regional cortical thickness, *X*) and behavioural (lifestyle risk factors linked to adverse aging, *Y*) data from 28 804 UK Biobank participants (for more details, see *Supplementary Materials – Section S4*). As in our simulated data analysis, we performed PLS on subsamples drawn from the broader pool of participants, asking if the significance, strength, and stability of different LVs depended on sample size. Informed by our results, we performed a final PLS analysis on the full set of 28 804 participants and explored which outcome metrics could discriminate which LVs may be of interest.

#### 3.2.1. Analysis 1: Sample size

Drawing 100 random subsamples at each of 10 sample sizes from our pool of UK Biobank participants, we saw that the number of significant brain-behaviour latent variables increased with *N* (*Fig. 3*). In this regard, the unrotated test was the most liberal method (passing nearly every LV of a possible 17 when *N* was large, on average), while rotating the behavioural component was the most conservative. While unrotated tests tended to consider late LVs significant in large samples, the split-half stability and covariance explained of these effects tended to be relatively low. Sample plots showing these trends across LVs 1, 3, 5, and 10 are shown in *Supplementary Fig. S10*.

**Figure 3:**
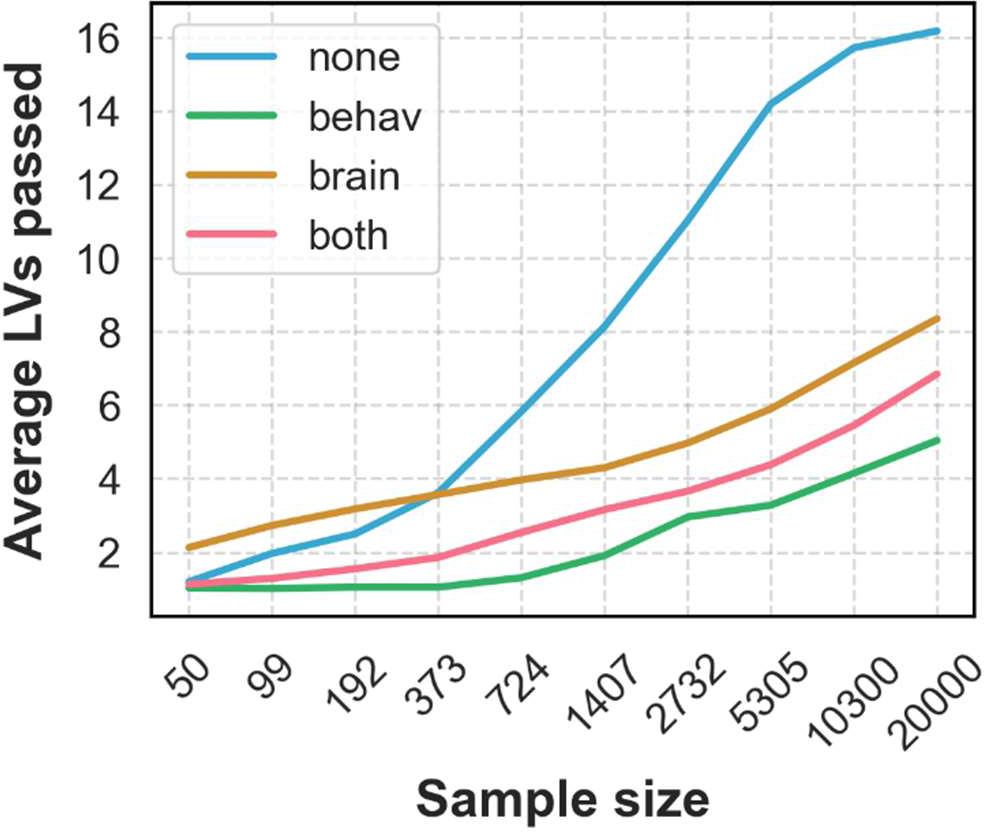
The average number of latent variables passing each permutation test (*p* < 0.05) across the 100 subsamples at each sample size. While the number of significant LVs increased with *N* for each method, the effect was especially pronounced for the unrotated tests, which passed nearly every possible LV in large samples.

#### 3.2.2. Analysis 2: Full sample

In our full sample of 28 804 participants, all permutation tests agreed that the first 5 LVs were significant (*Fig. 4*). However, the methods disagreed for further latent variables, with the unrotated test determining that *all* possible LVs were significant.

**Figure 4:**
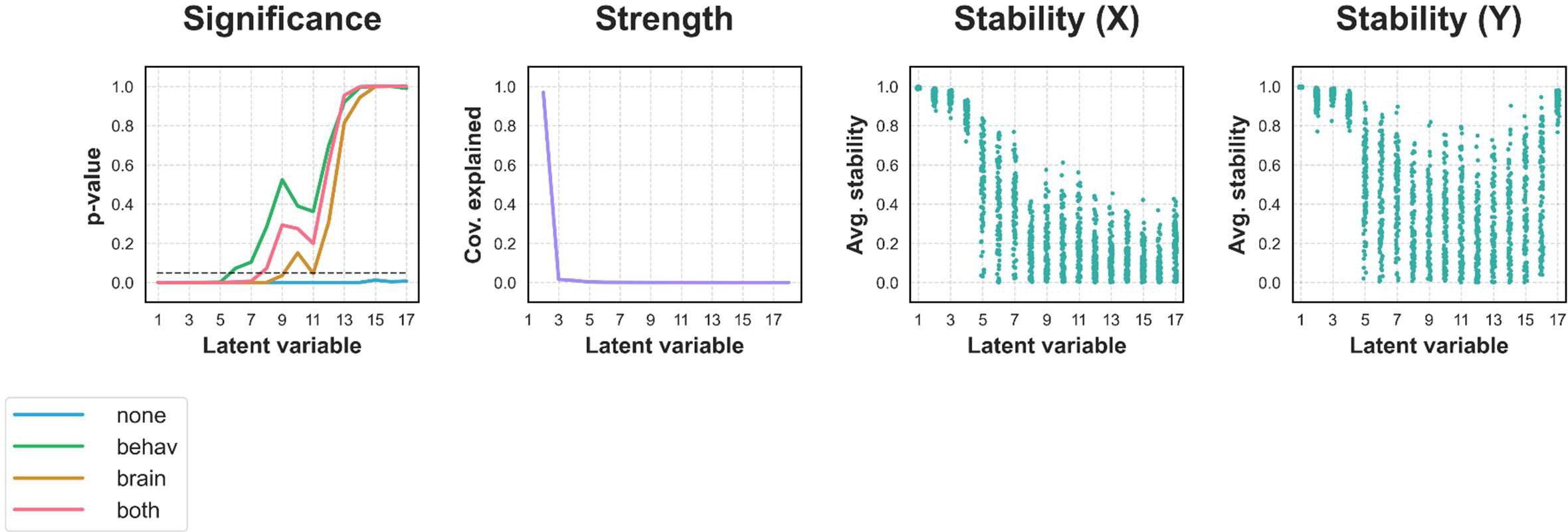
Latent variable metrics in the full UK Biobank sample. Regarding **Significance**, the unrotated test deemed every LV significant, while the rotated tests were far more conservative. Conversely, the **Strength** (covariance explained) of each LV dropped sharply after LV1, and the **Stability** of the brain **(X)** and behaviour **(Y)** LV components decreased and became far more variable after LV4 (individual points represent values for unique dataset splits).

While the unrotated test was entirely non-selective, our results in simulated data suggested that the error rates of rotated tests are unacceptably high for early LVs. Owing to the limitations of both approaches, we instead explore whether latent variable strength and stability metrics, which tracked key properties of our simulated data, can offer additional information regarding which effects may be relevant.

For one, split-half stability became far more variable across splits after LV4 (*Fig. 4*). In this case, a researcher may choose to set a stability threshold and report the first 4 LVs accordingly. Beyond being relatively stable, the first 4 LVs were also relatively interpretable. LV1 reflected globally thicker cortex in younger participants, while LV2, LV3, and LV4 reflected more focal cortical thickness patterns related most strongly to alcohol consumption, body-mass index, and education/smoking, respectively (*Fig. 5*). Behavioural loadings for subsequent LVs tended to be relatively high for variables which already loaded strongly onto previous LVs, or tended to be relatively scattered (*Supplementary Fig. S11*). In turn, a researcher may choose to report the first 4 latent variables, owing to their relative stability *and* coherence.

**Figure 5:**
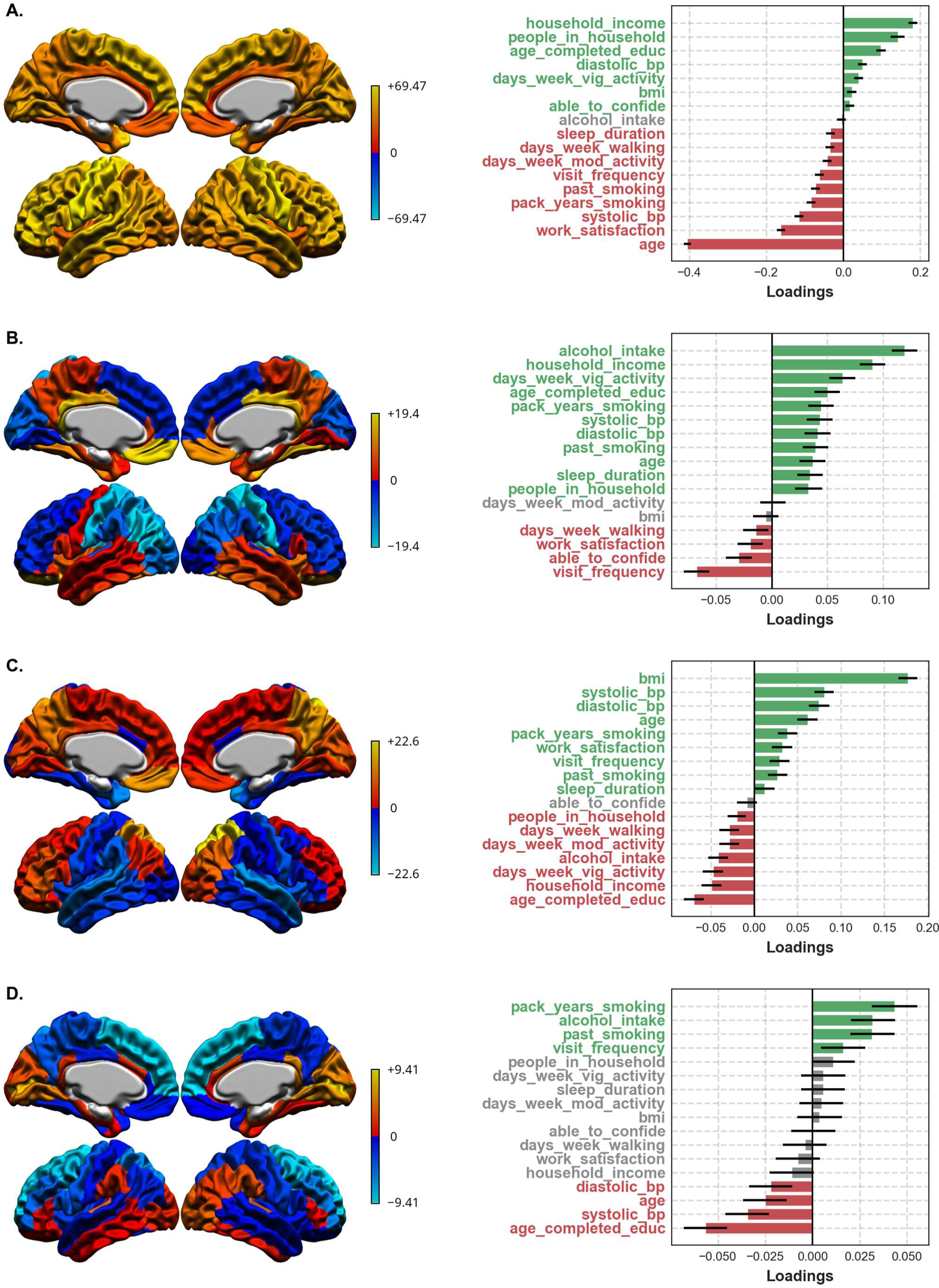
LVs 1-4 in the full UK Biobank sample. The first column shows bootstrap ratios for the brain variables, while the second shows loadings for the behavioural variables. **A. (LV1)** An expected pattern of broadly thicker cortex in younger participants. **B. (LV2)** A pattern of increases and decreases in cortical thickness across frontal, parietal, and cingulate regions related to greater alcohol consumption. **C. (LV3)** Broadly thicker cortex in the frontal and parietal lobes related to greater body-mass index. **D. (LV4)** Thinner superior frontal cortex related to greater smoking and lower education.

As a final example, a researcher may instead choose to only report the first latent variable, as it explained most of the covariance in the sample (*Fig. 4*). This may especially be the case if their research question is age-related, as LV1 is the dominant age-related pattern in the dataset, and all subsequent latent variables must capture covariance independent of it (see *Methods*).

## 4. Discussion

In a PLS analysis, latent variables are considered statistically significant if they pass permutation testing, with a Procrustes rotation applied to realign each set of permuted latent variables. However, it has not been shown whether applying the rotation, or applying the rotation to different components of the decomposition, provides a more accurate measure of whether true latent variables are present in a dataset, relative to the unrotated condition. Further, given growing concerns that the results from PLS and other multivariate techniques can fail to reproduce in held-out samples (Churchill et al., 2013; Nakua et al., 2024; Ji et al., 2021; Helmer et al., 2024; Dinga et al., 2019), it is plausible that significance testing can be complemented by measures of latent variable strength and stability to more thoroughly evaluate a PLS decomposition.

Accordingly, across a range of simulated datasets with a known ground truth, we observed that rotated permutation tests were characterized by high error rates for the first latent variable in randomly generated data, and routinely detected a simulated effect regardless of whether it was undersampled, weak, or noisy. Unrotated tests were more conservative in simulated data, but proved to be non-selective in large samples drawn from the UK Biobank.

Despite the limitations of the permutation tests, strength and stability metrics were relatively consistent: they accurately tracked our confidence that latent effects were present in simulated data, and showed potential for discriminating effects of interest in the UK Biobank.

The tendency for rotated tests to favour passing LV1 agrees with previous observations (Kovacevic et al., 2013) and follows from our understanding of PLS. Concretely, on a given permutation instance, the rows of one matrix are shuffled and PLS is performed under the null hypothesis of no cross-modality latent effects. The Procrustes rotation is applied to redistribute covariance explained across the permuted LVs (see *Methods –* *Eq. (7)*), shifting the null distributions which the original LVs are tested against. Critically, since LV1 is always the *strongest possible* source of covariance in a PLS decomposition, there is no situation where the permuted LV1 can “gain” covariance after the Procrustes rotation, as it was already as strong as possible beforehand. As such, following the rotation, covariance must tend to shift from early to late null distributions, making it easier to detect the strongest effect, and more difficult to detect weaker ones. For visual examples of this effect, where the expected decline in null distribution strengths becomes less pronounced following any rotation in both randomly generated and UK Biobank data, see *Supplementary Figs. S9* and *S12*. See also *Supplementary Materials – Section S1* for a mathematical description of how the Procrustes rotation systematically alters the LV1 null distribution.

As our results appear to be a consequence of the rotation itself, we expect rotated permutation tests to systematically favour passing LV1 in any application of PLS beyond analyzing brain-behaviour covariance. Further, the tendency for the Procrustes rotation to “dilute” the first of a set of magnitude-ordered effects should extend to other use cases of the technique beyond PLS permutation testing, including recent gradient mapping studies, where the rotation is used to align axes describing continuous cortical variability in an individual to those from a group average (Langs et al., 2015; Vos de Wael et al., 2020).

More generally, it is unclear whether applying the rotation during permutation testing has a strong theoretical backing. For one, the covariance explained by LV1 in any sample should overestimate that of the population (Lawley, 1956). If the strength of LV1 is already overestimated, and if rotated permutation tests make LV1 easier to detect, then the procedure may be suboptimal for PLS statistical inference. Further, the Procrustes rotation was originally brought to PLS and related techniques in the context of bootstrap resampling, not permutation testing (Milan & Whittaker, 1995). Relative to the original sample, any bootstrapped sample should contain the same latent effects, though they may be reordered due to sampling variability – justifying a transformation to map them to the originals (Milan & Whittaker, 1995). However, it is unclear whether such a situation holds for the permutation test, where latent effects are not simply reordered, but fundamentally *broken* by the reshuffling procedure. Finally, it should be noted that there is active research in the broader field of Procrustes analysis, which studies the optimal means of mapping one matrix to another. The orthogonal Procrustes problem, solved in PLS permutation testing, is just one way to perform a Procrustes analysis, and several other methods could equally be employed (a detailed discussion is beyond the scope of this report – see Gower (2010), Meng et al. (2022) for more details).

Beyond the permutation testing approach considered here, an alternative technique was recently proposed by Nakua et al. (2024). Briefly, the test considers whether the total covariance explained by a *set* of latent variables is greater than expected by chance (i.e.., the sum of the covariance captured by LV2-LV*N* must be significant for LV2 to be considered significant). This approach is similar to permutation inference based on the Wilk’s λ statistic, commonly used for canonical correlation analysis, a sister technique of PLS (Winkler et al., 2020). Importantly, the approach both avoids applying a Procrustes rotation and is thought to perform well in large samples (Nakua et al., 2024), obviating the concerns raised in this report about unrotated tests in large samples and rotated tests more generally.

Alongside permutation testing, we also assessed whether latent variable strength and stability were sensitive to key properties of simulated data. Tellingly, the split-half stability and covariance explained of LV1 tended to be relatively low when the simulated effect was undersampled, weak, or noisy, and when PLS was performed on whitened data. Our findings agree with early observations stressing the complementary nature of significance and stability in a PLS analysis (McIntosh & Lobaugh, 2004), as well as recent simulation studies, which have reported that stability metrics were sensitive to sample size (Helmer et al., 2024), effect strength (Kovacevic et al., 2013), noise (Kovacevic et al., 2013), and whether an effect was encoded (McIntosh, 2022). Accordingly, beyond PLS, our results add to a growing body of literature showing that significant effects are not necessarily strong or stable across samples, stressing the importance of incorporating statistical analyses beyond significance testing, especially in multivariate paradigms (Chen et al., 2017; Dinga et al., 2019).

While we demonstrated that latent variable strength and stability metrics were coupled to a simulated ground truth, we did not provide a framework based on these measures for selecting which latent variables may be of interest. However, many such frameworks have been described across the PLS literature. For one, Carbonell (2020) simply reported LV1, which by definition captured the most covariance in the sample (regardless, one could argue that any effects after LV1 are somewhat artificial, as they are all constrained to be independent of it). Meanwhile, Misic et al. (2016) analyzed LVs which passed two tests for covariance explained commonly seen in the principal components analysis literature. Specifically, they tested whether an LV was stronger than the average LV in the dataset, and whether it passed a visual scree test for covariance explained. Regarding split-half stability, McIntosh (2022) and Nakua et al. (2024) calculated *z*-values describing the distributions of stability values for each LV, and suggested that LVs with a *z*-value above a heuristic threshold can be considered consistently stable.

Finally, much like assessing stability, Hansen et al. (2021), Kirschner et al. (2020), and Mirchi et al. (2019) all cross-validated their PLS results by assessing whether the cross-modality relationships captured by LVs in a training dataset were sufficiently strong in a held-out sample. Based on our findings, all such approaches may serve as reasonable alternatives or complements to conventional permutation testing.

Together, rotated permutation tests are systematically more permissive of early latent variables, while unrotated permutation tests can be exceedingly liberal in large samples. Owing to the limitations of both approaches, we argue that the *p*-values from either test cannot be interpreted as strict measures of whether a latent variable “exists”, and that tests based on latent variable strength and stability should be considered as part of a more nuanced formula for determining the effects to report in a PLS analysis. We end by presenting a set of considerations based on our findings for researchers implementing PLS permutation testing, shown in *Table 1*.

**Table 1:**
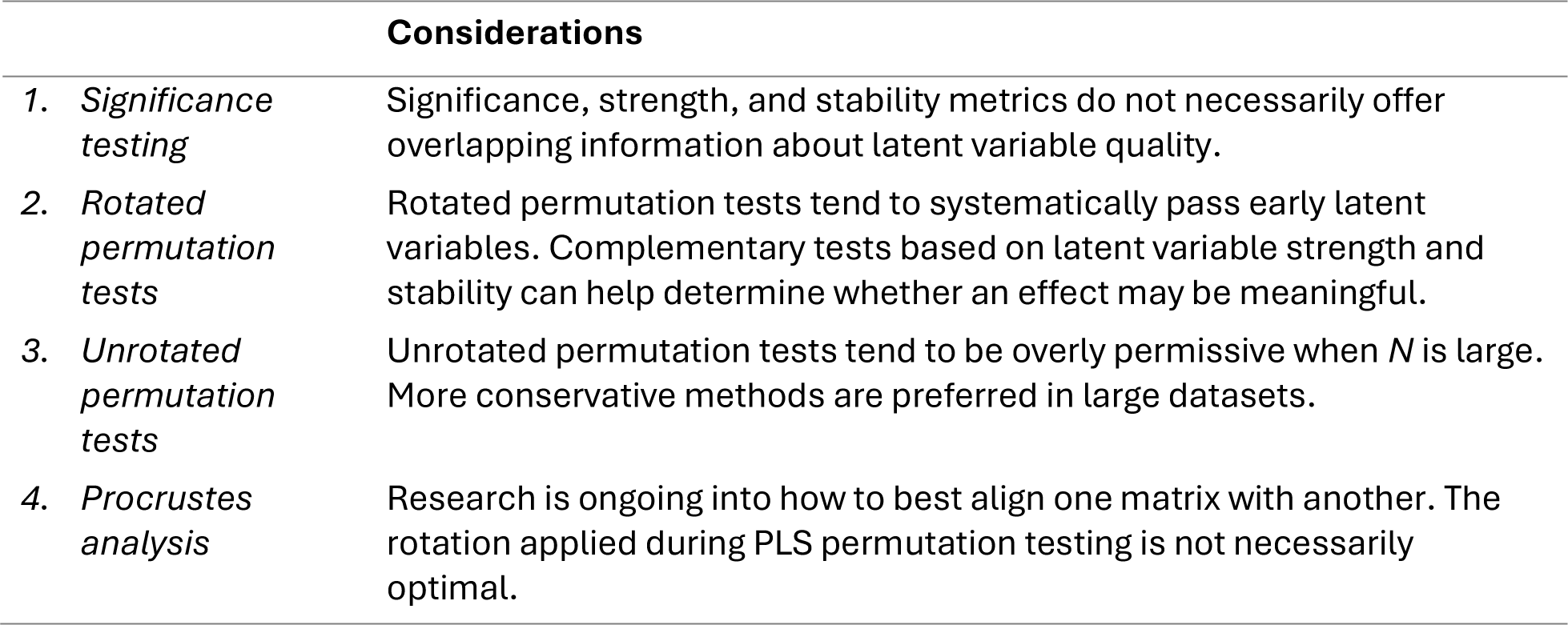
Considerations for PLS permutation testing.

## Data and Code Availability

The code used to reproducibly generate the simulated datasets and perform the data analysis is available on GitHub (https://github.com/danyluik/pls_rotation_analysis).

## Author Contributions

**MD:** Conceptualization, Software, Formal Analysis, Investigation, Writing – Original Draft, Writing – Review & Editing. **YZ:** Conceptualization, Formal Analysis, Writing – Review & Editing. **AM:** Data Curation. **ML:** Data Curation. **JS:** Data Curation. **RJ:** Data Curation. **BM:** Conceptualization, Writing – Review & Editing. **YIM:** Conceptualization, Supervision, Writing – Review & Editing. **MMC:** Conceptualization, Supervision, Writing – Review & Editing, Funding Acquisition.

## Acknowledgements

The authors thank T. Robert Baumeister for helpful initial conversations.

