## supplemental for "Evaluating permutation-based inference for partial least squares analysis of neuroimaging data"

### S. Supplementary Materials

#### S1. Impact of the Procrustes rotation on PLS null distributions

Below, we consider how the Procrustes rotation systematically alters the LV1 null distribution. As described in the *Methods*, assume we rotate the right singular vectors on a given permutation instance. Of relevance, we have permuted singular vectors $V^{(p)}$, permuted singular values $\Sigma^{(p)}$, and the Procrustes rotation matrix $R$ which, when applied to $V^{(p)}$, aligns it as closely as possible with the right singular vectors from the original decomposition, $V$. Assuming no rotation is applied, the values for each null distribution are simply given by the diagonal of $\Sigma^{(p)}$. However, following the Procrustes rotation, the adjusted null distribution values are given by:

$$\tilde{V}^{(p)} = V^{(p)}\Sigma^{(p)}R \tag{1}$$

$$val_i = ||(\tilde{V}^{(p)})_i||_2 \tag{2}$$

Where *i* represents a column of $\tilde{V}^{(p)}$. Given that the L2 norm is sub-multiplicative:

$$val_i \leq ||(V^{(p)})_i||_2 \, ||(\Sigma^{(p)}R)_i||_2 \tag{3}$$

Since $V^{(p)}$ contains singular vectors, $||(V^{(p)})_i||_2 = 1$, and *Eq. (3)* can equivalently be expressed as:

$$val_i \leq ||(\Sigma^{(p)}R)_i||_2 \tag{4}$$

Which can be expanded to:

$$val_i \leq \left\| R_{1i}\sigma_1^{(p)} + R_{2i}\sigma_2^{(p)} + \cdots + R_{ni}\sigma_n^{(p)} \right\|_2 \tag{5}$$

Where *n* represents the number of singular vectors. Before proceeding, we note that $\Sigma^{(p)}$ contains singular values along its diagonal, such that:

$$\sigma_1^{(p)} > \sigma_2^{(p)} > \cdots > \sigma_n^{(p)} \tag{6}$$

And that *R* is orthonormal (Schönemann, 1966), such that:

$$||R_i||_2 = 1 \; \forall i \tag{7}$$

Given that the upper limit of the L2 norm of a vector is the sum of its elements (i.e., the L1 norm), the sum of the individual elements of $R_i$ must not exceed 1. Therefore, the right side of *Eq. (5)* can be seen as the L2 norm of a set of weighted singular values, where the sum of the weights cannot exceed 1. Next, we consider the case where $i = 1$:

$$val_1 \leq \left\| R_{11}\sigma_1^{(p)} + R_{21}\sigma_2^{(p)} + \cdots + R_{n1}\sigma_n^{(p)} \right\|_2 \tag{8}$$

Following *Eq. (6)* and *(7)*, we can infer that:

$$R_{11}\sigma_1^{(p)} + R_{21}\sigma_2^{(p)} + \cdots + R_{n1}\sigma_n^{(p)} \leq (R_{11} + R_{21} + \cdots + R_{n1})\sigma_1^{(p)} \leq \sigma_1^{(p)} \tag{9}$$

Again, given that the upper limit of the L2 norm of a vector is the sum of its elements:

$$\left\| R_{11}\sigma_1^{(p)} + R_{21}\sigma_2^{(p)} + \cdots + R_{n1}\sigma_n^{(p)} \right\|_2 \leq \sigma_1^{(p)} \tag{10}$$

Substituting *Eq. (10)* into *Eq. (8)*, we can conclude:

$$val_1 \leq \sigma_1^{(p)} \tag{11}$$

As such, following a Procrustes rotation on a given permutation instance, the value which contributes to the LV1 null distribution must be less than or equal to that from the unrotated condition.

#### S2. Preliminary findings

Data were collected through the recruitment of clients and their families at the *Prevention and Early Intervention for Psychosis* and *Clinic for Assessment of Youth at Risk* services at the Douglas Mental Health University Institute in Montreal, Canada (Pruessner et al., 2017). Participants were either diagnosed with a first episode of psychosis within 3 months of study participation, at clinical high-risk for psychosis (expressing subthreshold symptoms according to CAARMS criteria (Pruessner et al., 2017)), or at familial high-risk for psychosis (siblings of the psychosis patients in the study). All participants provided written informed consent for the secondary analysis of their data, and the study was approved by a Research Ethics Board at the Douglas Mental Health University Institute.

All images were recorded with a 3T Simens Magnetom scanner. T1-weighted images (MPRAGE, 1 $mm^3$) were collected for each participant and preprocessed using the minc-bpipe-library (https://github.com/CoBrALab/minc-bpipe-library). Images were then processed with CIVET 2.1.1 (Kim et al., 2005), giving cortical thickness at 77 122 vertices across the cortical surface. Positive and negative psychotic symptoms were measured using the 9 global items from the interviewer-assessed Scales for the Assessment of Positive/Negative Symptoms, and cognitive performance was summarized across the 7 domains of the CogState computerized research battery (Benoit et al., 2015).

Our final sample included 125 participants whose images passed strict motion (https://github.com/CoBrALab/documentation/wiki/Motion-Quality-Control-(QC)-Manual) and CIVET (https://github.com/CoBrALab/documentation/wiki/CIVET-Quality-Control-Guidelines) quality control guidelines and whose symptom measures were complete. Accordingly, for our PLS analysis, the brain matrix was composed of $(125 \cdot 77\,122)$ vertex-wise cortical thickness measures and the behavioural matrix was composed of $(125 \cdot 16)$ clinical and cognitive measures. PLS was performed as described in *Section 2.3.5* with the 4 permutation test methods. To confirm that our results were not driven by an inadequate number of permutations, the input matrices were reshuffled 100 000 times.

The *p*-values by LV for each permutation test are shown in *Supplementary Fig. S1*. All tests agreed that LV1 was significant, but the *p*-values for LV2 dramatically diverged, ranging from significant when rotating the brain component to approaching 1 when no rotation was applied. For reference, the LV2 null distributions are shown in *Supplementary Fig. S2*.

As LV2 was relatively coherent, linking focal cortical thickness reductions to a clear pattern of positive and disorganized psychotic symptoms (*Supplementary Fig. S3*), we sought to systematically evaluate which permutation test should be “trusted” regarding its significance, motivating the present study.

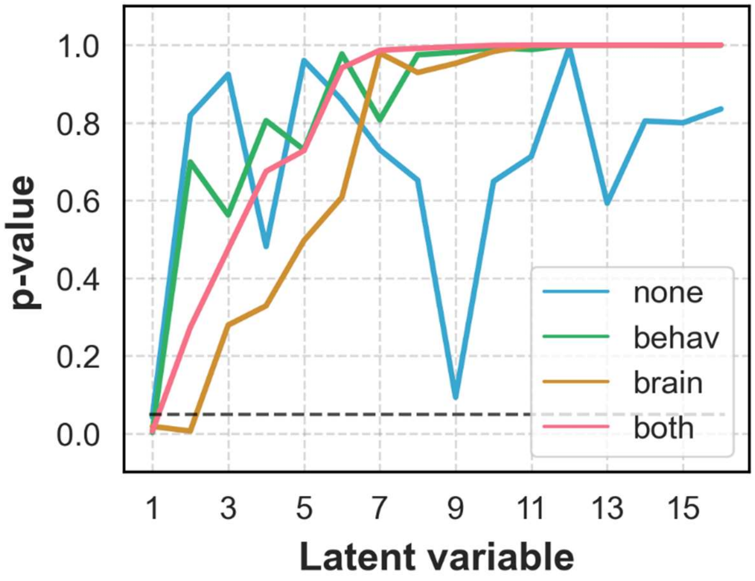


**Figure S1**: *p*-values by latent variable for each permutation test performed in our psychosis spectrum dataset. The dashed line indicates a conventional significance threshold ($p < 0.05$). *p*-values began to meaningfully diverge after LV1.

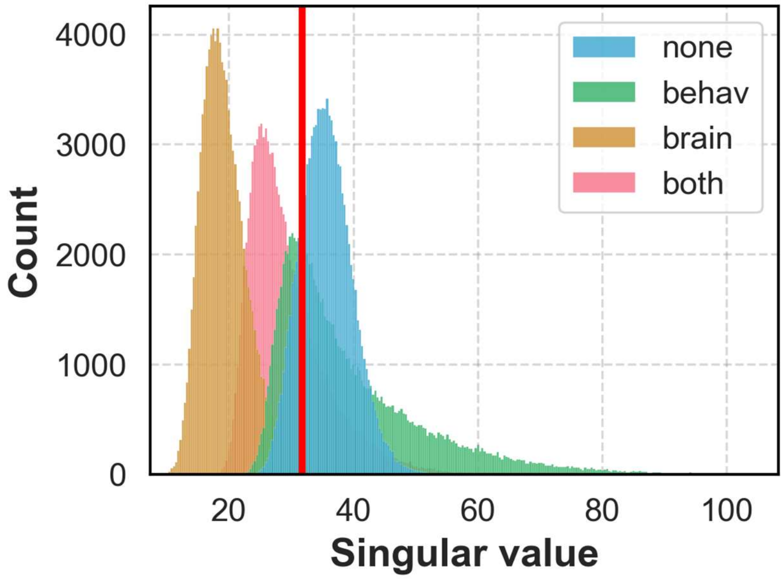


**Figure S2:** LV2 null distributions for each permutation test performed in our psychosis spectrum dataset. The red line represents the LV2 singular value from the initial PLS decomposition.

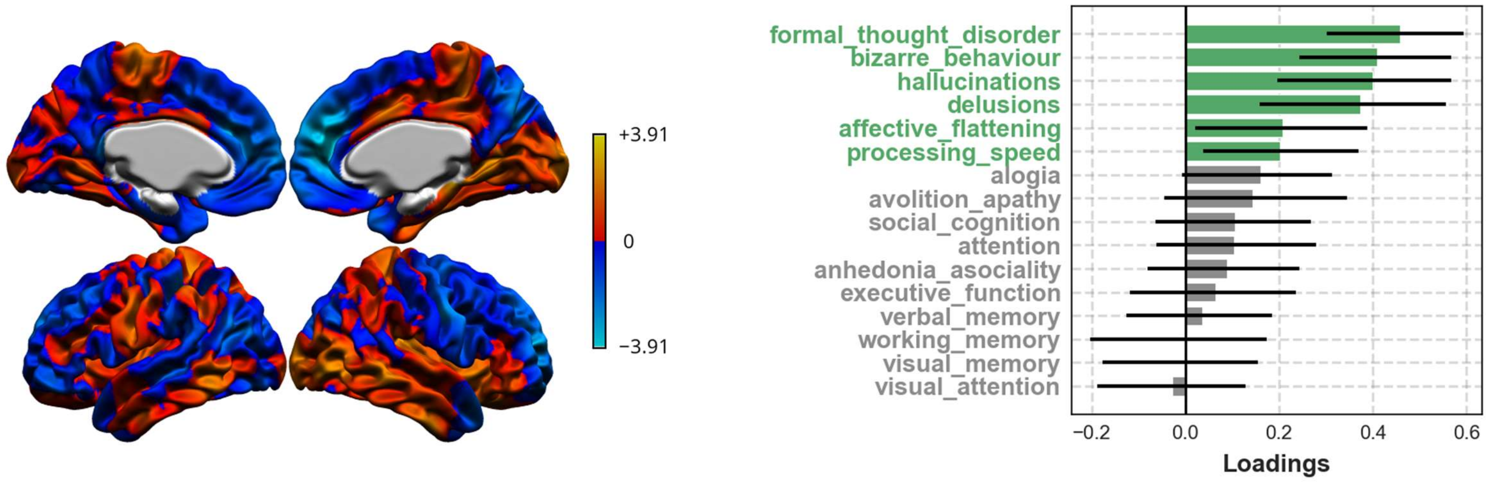


**Figure S3:** LV2 in our psychosis spectrum sample, showing bootstrap ratios for brain variables and loadings for behavioural variables, respectively. Combined, the LV mainly reflects thinner prefrontal cortex related to positive and disorganized symptoms.

#### S3. Simulated data

##### *S3.1. Features*

We explored whether our PLS outcome metrics were related to the input dimensionality. Simulating data using GEMMR, we increased the number of features in the $Y$ matrix in 10 steps logarithmically spaced between 2 and 200, with 100 different datasets per dimensionality. The number of latent effects ($m = 1$), canonical correlation ($r = 0.3$), and sample size ($N = 1000$) were held constant.

After performing PLS on each dataset, we observed that all permutation tests consistently detected LV1 regardless of the number of features (*Supplementary Fig. S4A*). LV1 strength and the stability of the $Y$ matrix declined monotonically with the dimensionality, while the stability of $X$ remained relatively constant (*Supplementary Fig. S4A*).

##### *S3.2. Whitened data*

We also wondered whether PLS permutation tests could “find” latent effects in datasets where the features were constrained to not meaningfully covary. Accordingly, we generated $X$ and $Y$ matrices of random values drawn from a normal distribution with mean 0 and variance 1. Then, we standardized the columns of either matrix to have mean 0 and variance 1 and whitened the result by rotating it to its principal component space. Using this procedure, we generated 100 datasets at each of 10 sample sizes, with sample sizes log-spaced between 100 and 10 000. The number of features (90 $X$, 10 $Y$) was held constant across datasets. Note that our simulation protocol differs slightly from our other analyses where we tuned sample size (*Fig. 2A-B*). The lowest sample size was set to 100 rather than 10, ensuring that each matrix always had more observations than features, and that it could be decorrelated in turn.

We assessed the efficacy of our whitening procedure by quantifying the distance of each covariance matrix from the identity matrix as follows (with $C_{XX}$ as an example):

$$Error = ||C_{XX} - I||_F$$

We observed that errors negligibly diverged from 0 (*Supplementary Fig. S5*) and proceeded with our analysis in turn. Here, all rotated permutation tests considered LV1 significant regardless of *N*, while the unrotated tests rarely detected the unsimulated “effect” (*Supplementary Fig. S4B*). Meanwhile, LV1 strength and stability metrics increased marginally with sample size, though both sets of metrics plateaued at relatively low values (*Supplementary Fig. S4B*).

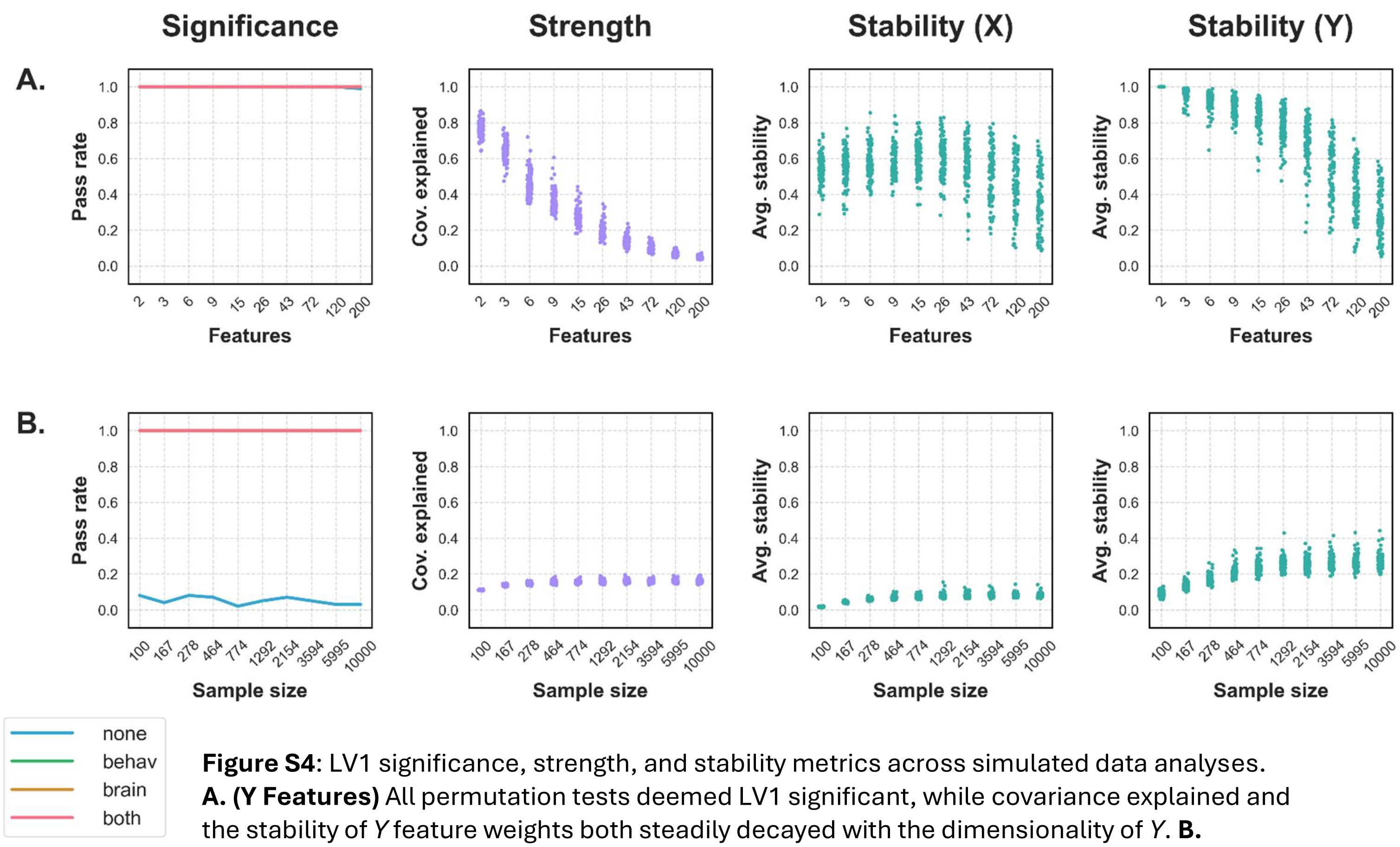


**Figure S4**: LV1 significance, strength, and stability metrics across simulated data analyses. **A. (Y Features)** All permutation tests deemed LV1 significant, while covariance explained and the stability of *Y* feature weights both steadily decayed with the dimensionality of *Y*. **B. (Whitened data)** Rotated tests consistently deemed LV1 significant, even though the data was constrained to not covary. Meanwhile, the unrotated test was less likely to detect an effect, and strength and stability metrics remained low regardless of sample size.

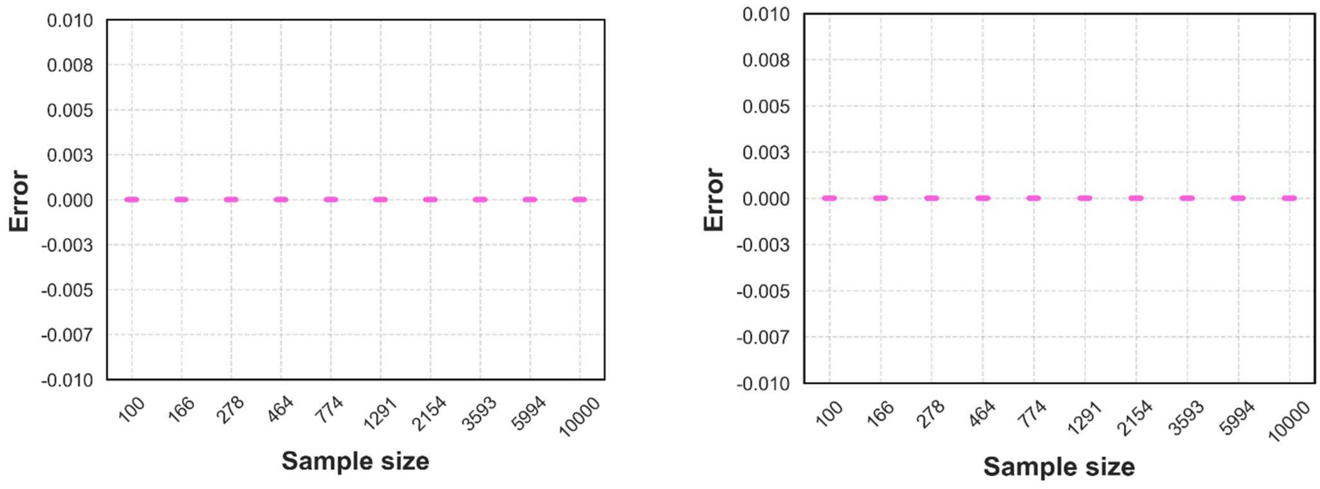


**Figure S5**: error (see *Section S3.2*) of whitening procedure for each *X* (left) and *Y* (right) matrix. Error values did not qualitatively differ from 0.

##### *S3.3. Z-scores*

As we binarized each permutation test into "pass" or "fail", we were also interested in more precisely evaluating how the rotations impacted our null distributions. Accordingly, for a given singular value $\sigma$ and its corresponding null distribution with mean $\bar{x}$ and standard deviation $s$, we calculated the z-value of $\sigma$ within the null distribution as:

$$z = \frac{\sigma - \bar{x}}{s}$$

Results for each supplemental analysis are shown *Supplementary Fig. S6*, and those from the main text are shown in *Supplementary Fig. S7*.

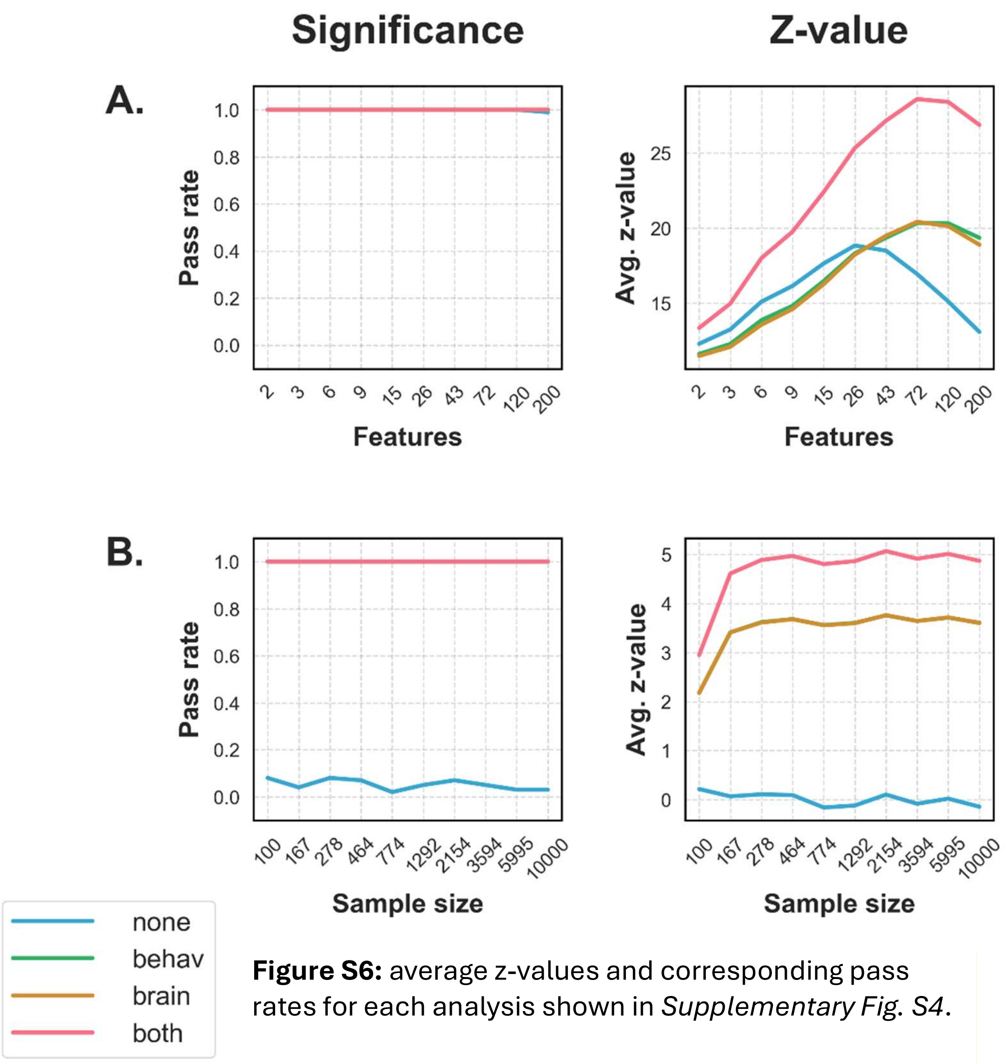


**Figure S6:** average z-values and corresponding pass rates for each analysis shown in *Supplementary Fig. S4*.

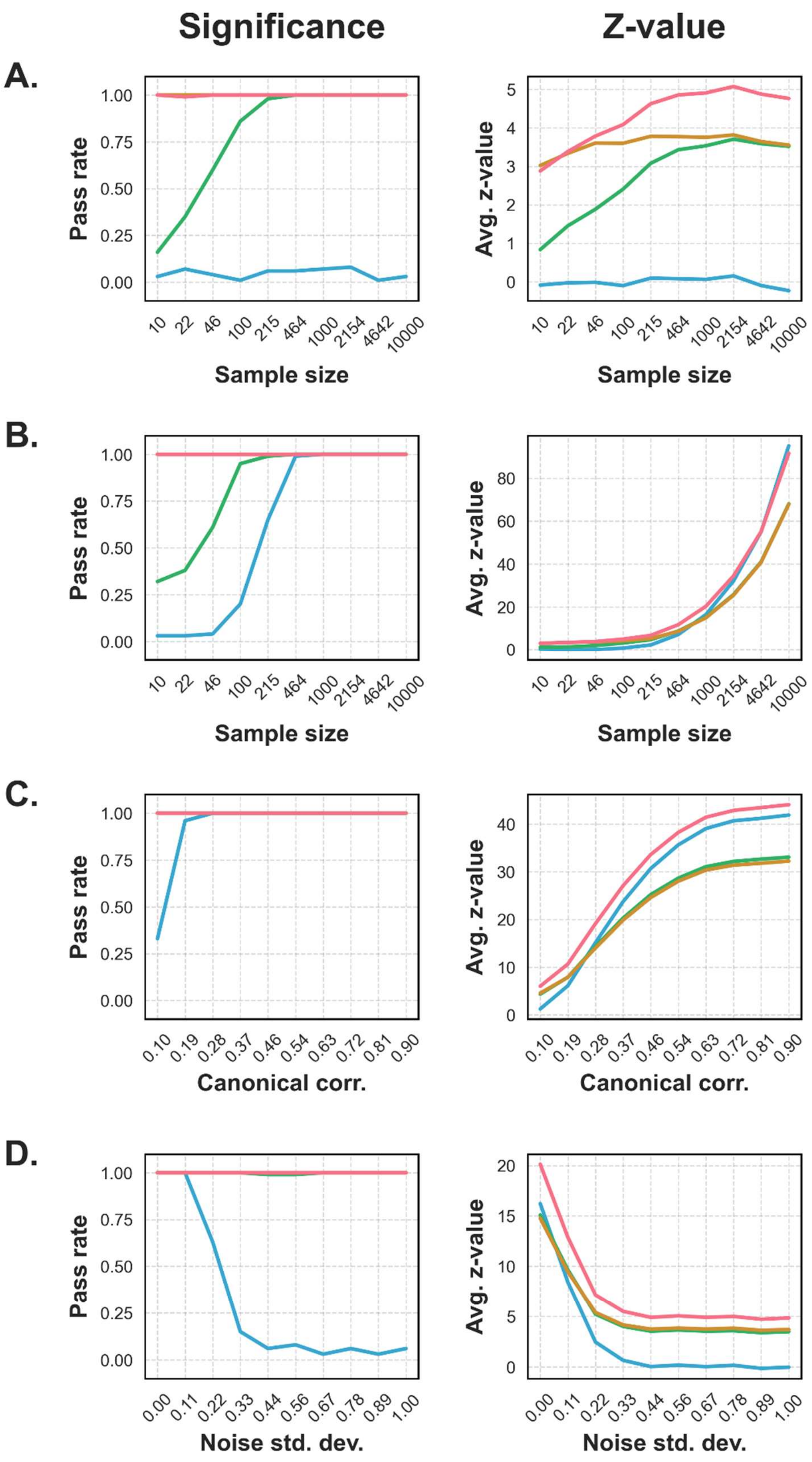


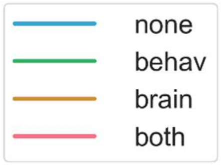


**Figure S7:** average z-values and corresponding pass rates for each analysis shown in *Fig. 2*.

##### *S3.4. Additional figures*

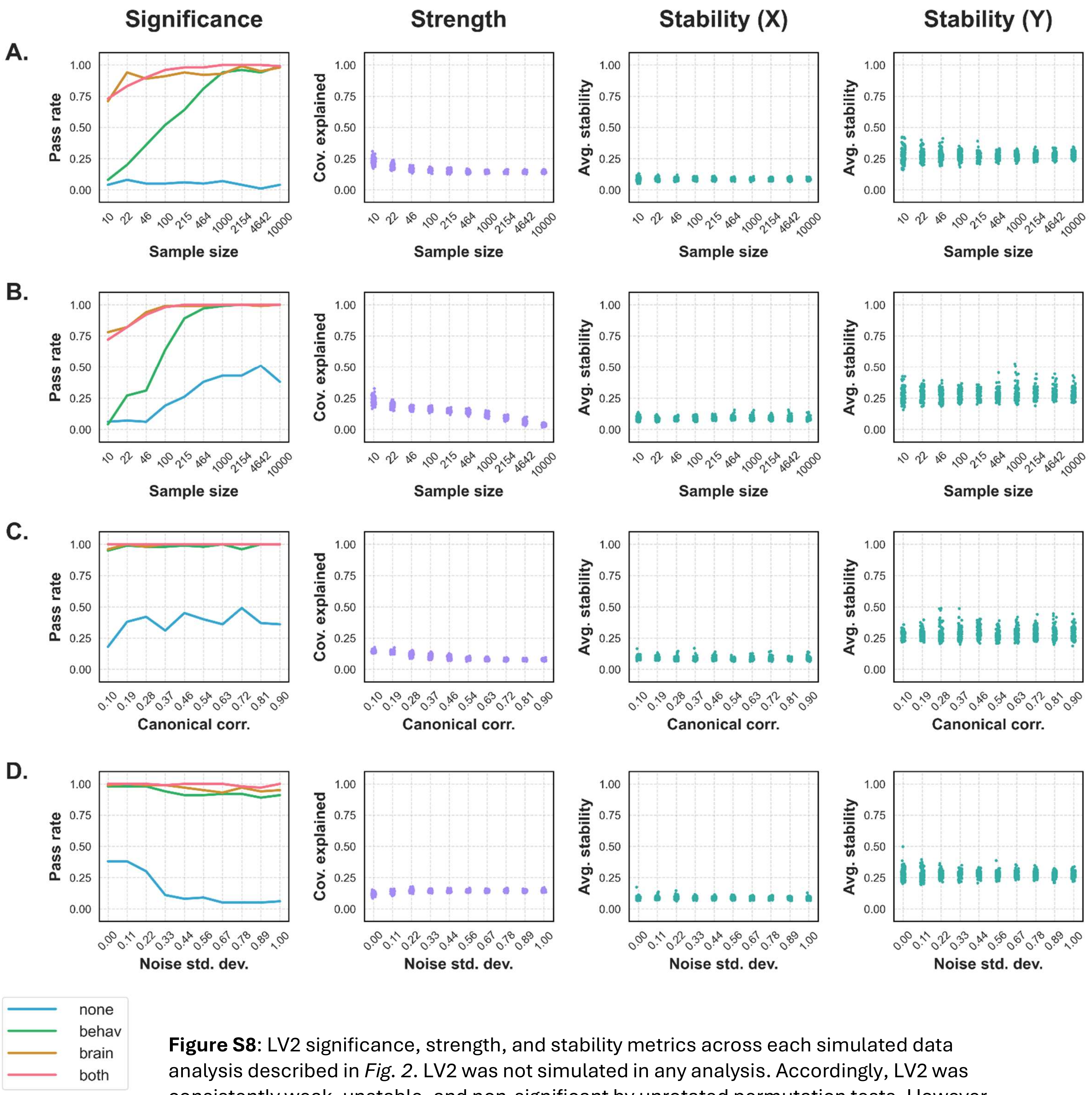


**Figure S8**: LV2 significance, strength, and stability metrics across each simulated data analysis described in *Fig. 2*. LV2 was not simulated in any analysis. Accordingly, LV2 was consistently weak, unstable, and non-significant by unrotated permutation tests. However, rotated permutation tests consistently detected LV2, suggesting that they are characterized by unacceptably high error rates.

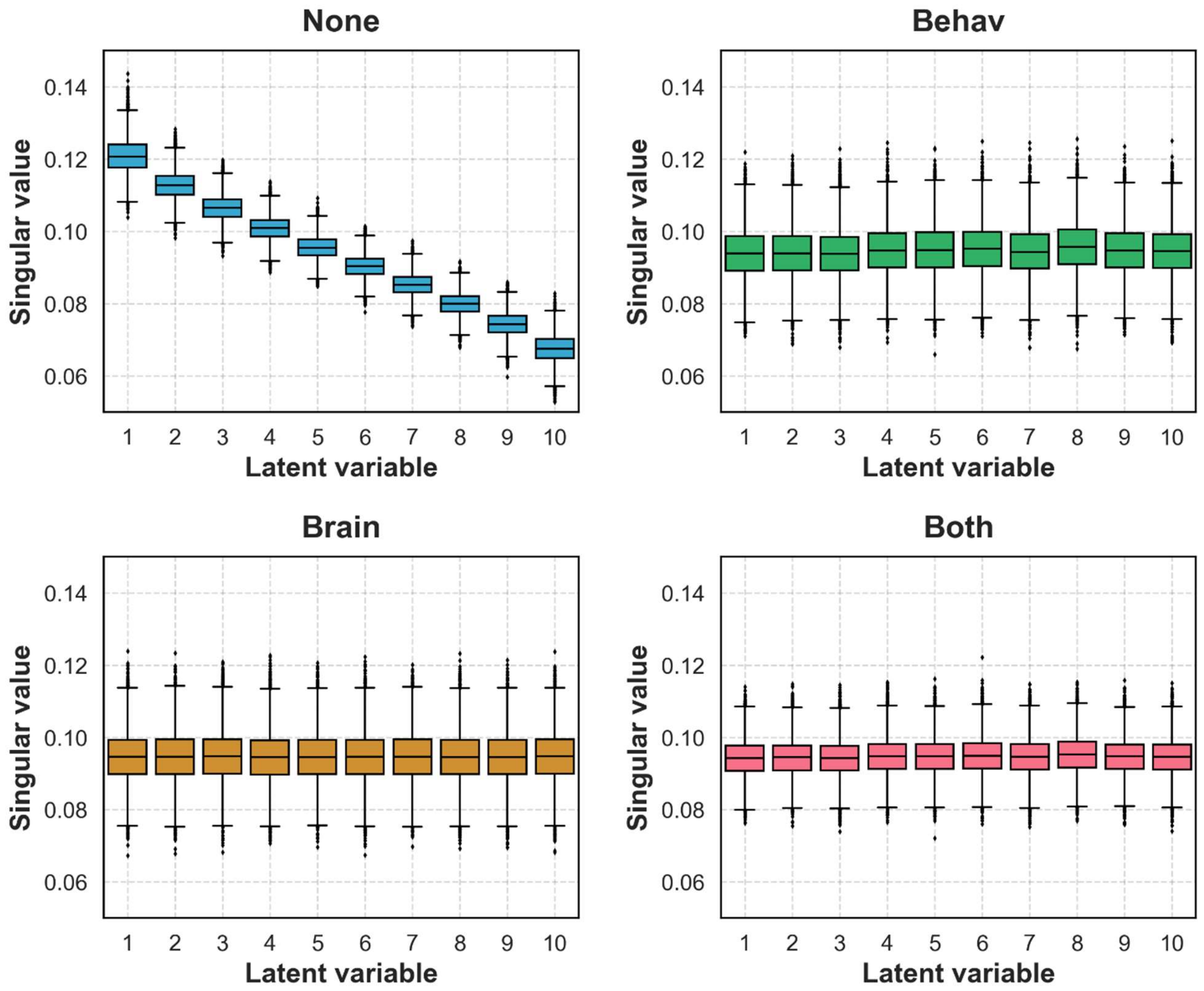


**Figure S9*:*** null distributions by latent variable for a randomly generated dataset (*N* = 10 000), plotted separately for each permutation test method. When no rotation was applied (top left), the means of the null distributions steadily decayed. However, after any rotation was applied, the null distributions became visually indistinguishable – an unreasonable null, given that covariance explained must decrease with LV in any PLS decomposition. By extension, when any rotation was applied, early LVs were tested against "weaker" null distributions and late LVs were tested against "stronger" ones, relative to the unrotated condition.

#### S4. UK Biobank

##### *S4.1. Sample*

Open access data were obtained from the UK Biobank, a population cohort of 500 000+ cognitively healthy participants aged 40 to 69 recruited across the United Kingdom between 2006 and 2010. All participants were registered with the National Health Service, lived within reasonable traveling distance of an assessment center, and provided informed written consent for the study. Full details of recruitment and other UK Biobank procedures are available online (http://www.ukbiobank.ac.uk/). Starting in 2014, a subset of 100 000 participants were invited to undergo multimodal magnetic resonance imaging (MRI). Of these individuals, we selected a cohort of 28 804 participants aged 45 to 82 at time of imaging who met the standards detailed below. Participants with a confounding neurological condition were excluded from the analysis.

##### *S4.2. Image processing*

Data were collected with a standard Siemens Skyra 3T. Comprehensive documentation on the imaging protocol can be found online (https://biobank.ctsu.ox.ac.uk/crystal/crystal/docs/bmri_V4_23092014.pdf) . We performed manual quality control as above for motion on T1-weighted (MPRAGE, 1 $mm^3$) scans prior to image processing. Scans passing quality control were input to CIVET 2.1.1 (Kim et al., 2005) to obtain regional average cortical thickness according to the Desikan-Killiany atlas.

##### *S4.3. Lifestyle variables*

Behavioral, demographic, and health-related data were collected from participants at each visit to the assessment center, and here, preference was given to answers provided during the imaging visit. 38 lifestyle risk factors were selected and analyzed for a separate study, and participants were excluded if they were missing measurements for at least 10 of these variables. All further missing data was imputed using the missForest package in R. For the present study, we selected a subset of 17 of these variables with variance of at least 1 across the sample. Each is listed and described below. Questionnaire details are available online (https://biobank.ndph.ox.ac.uk/showcase/).

- *Age*: at imaging visit.
- *Alcohol intake frequency*: provided on a 5-point scale, with answer options ranging from “Never” to “Daily or almost daily”.
- *Age of completing full time education*: coded as 21 for any participant who indicated having a tertiary degree (log-transformed).
- *Body-mass index*: calculated from participant height and weight measured at the assessment centre, in $kg/m^2$.
- *Days per week of walking 10+ minutes*
- *Days per week of moderate physical activity*
- *Days per week of vigorous physical activity*

- *Average total household income*: on a 5-point scale from “Less than 18 000” to “Greater than 100 000”, in £ before tax.
- *Sleep duration*: average, in hours.
- *Past tobacco smoking*: provided on a 4-point scale, ranging from “I have never smoked” to “Smoked on most or all days”.
- *Pack years of smoking:* number of cigarettes smoked per day, divided by 20, and multiplied by the number of years spent smoking (log-transformed).
- *Ability to confide:* provided on a 6-point scale, ranging from “Never or almost never” to “Daily or almost daily”.
- *Frequency of family/friend visits*: provided on a 6-point scale, ranging from “Never or almost never” to “Daily or almost daily”.
- *People in household*: provided on a 4-point scale, ranging from “Only me” to “Five or more”.
- *Work/job satisfaction*: provided on a 7-point scale, with the lowest value corresponding to “I am not employed”, and subsequent values ranging from “Extremely unhappy” to “Extremely happy”.
- *Systolic blood pressure*: measured manually or automatically by registered nurses using an Omron 705 IT electronic blood pressure monitor.
- *Diastolic blood pressure*: as above.

##### *S4.4. Additional figures*

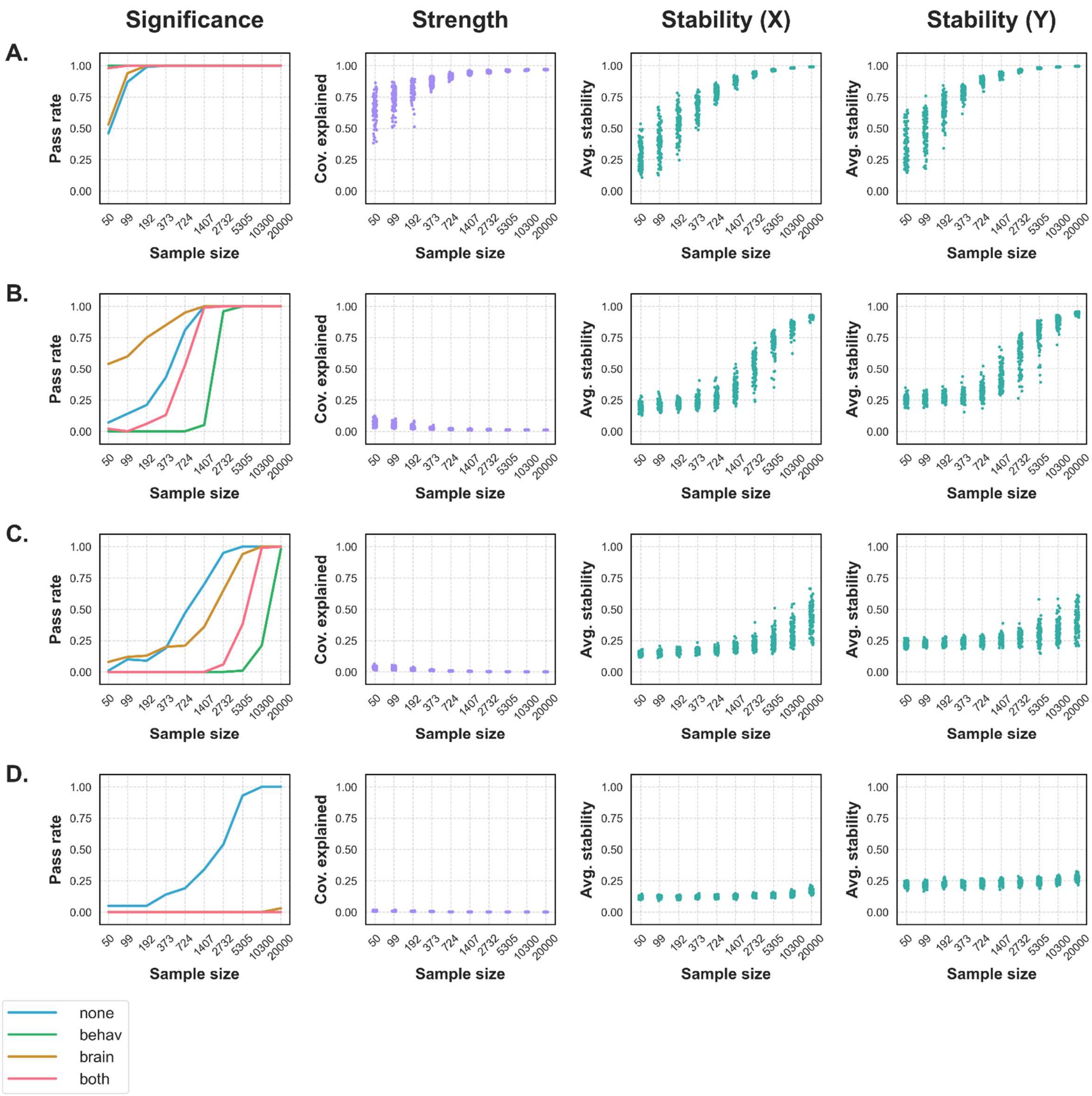

**Figure S10**: LV1, 3 5, and 10 significance, strength, and stability metrics across subsamples of our UK Biobank sample. Columns are described in *Fig. 2*. **A. (LV1)** LV1 was consistently considered significant, strong, and stable at most sample sizes. **B. (LV3)** The permutation tests disagreed regarding LV3 significance, except at the largest sample sizes. Notably, the unrotated permutation test is not the most conservative method. While LV3 did not explain much covariance, it was considered stable in large samples. **C. (LV5)** The permutation tests again disagreed on LV significance, with the unrotated test now the most liberal method. Meanwhile, LV5 was not consistently considered strong or stable. **D. (LV10)** Only the unrotated test considered LV10 significant, and only in the largest samples. LV10 was never considered strong or stable.

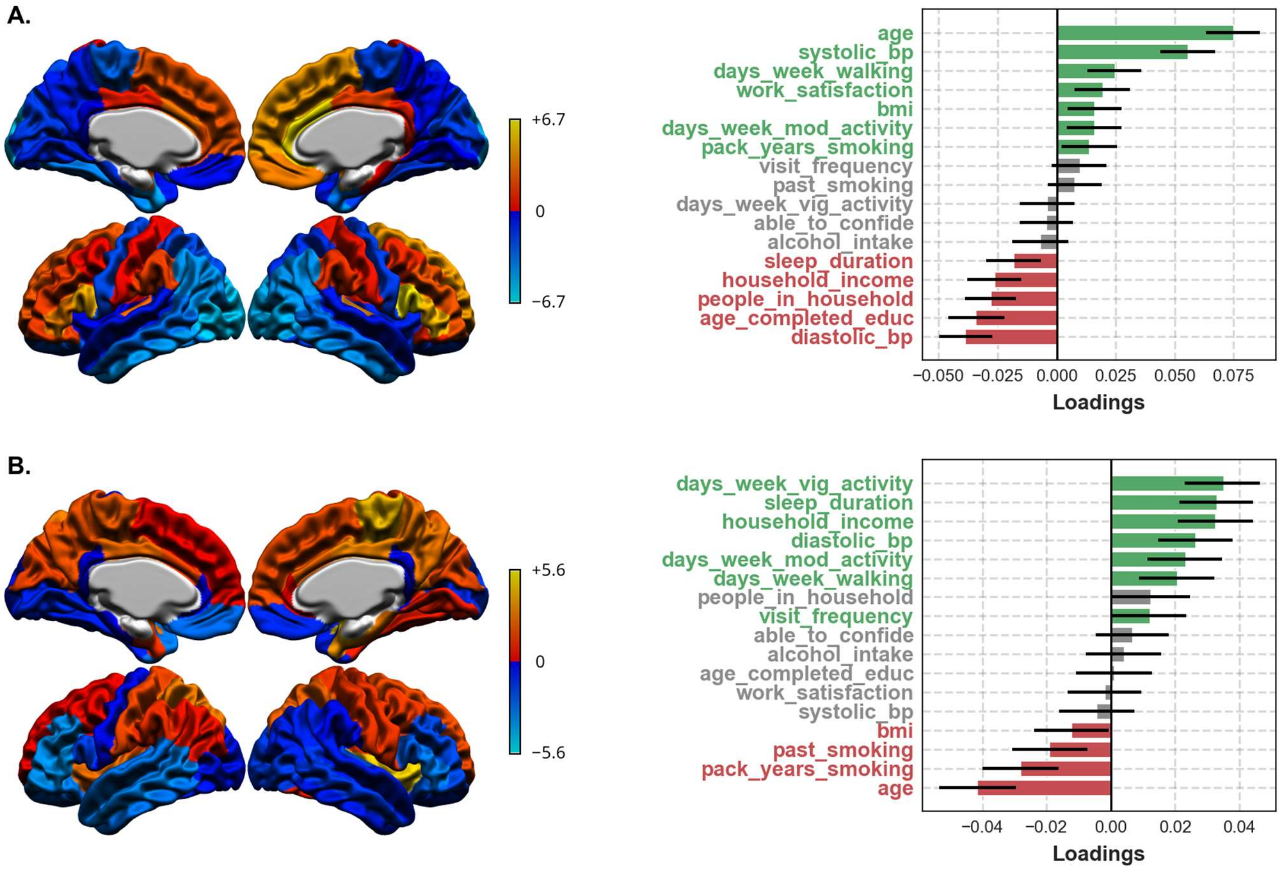


**Figure S11**: LVs 5-6 in the full UKBB sample. **A. (LV5)** Combined, the LV reflects a cortical thickness pattern related most strongly to age, which was already a dominant contributor to LV1. **B. (LV6)** Together, no dominant sets of brain or behavioural variables were observed.

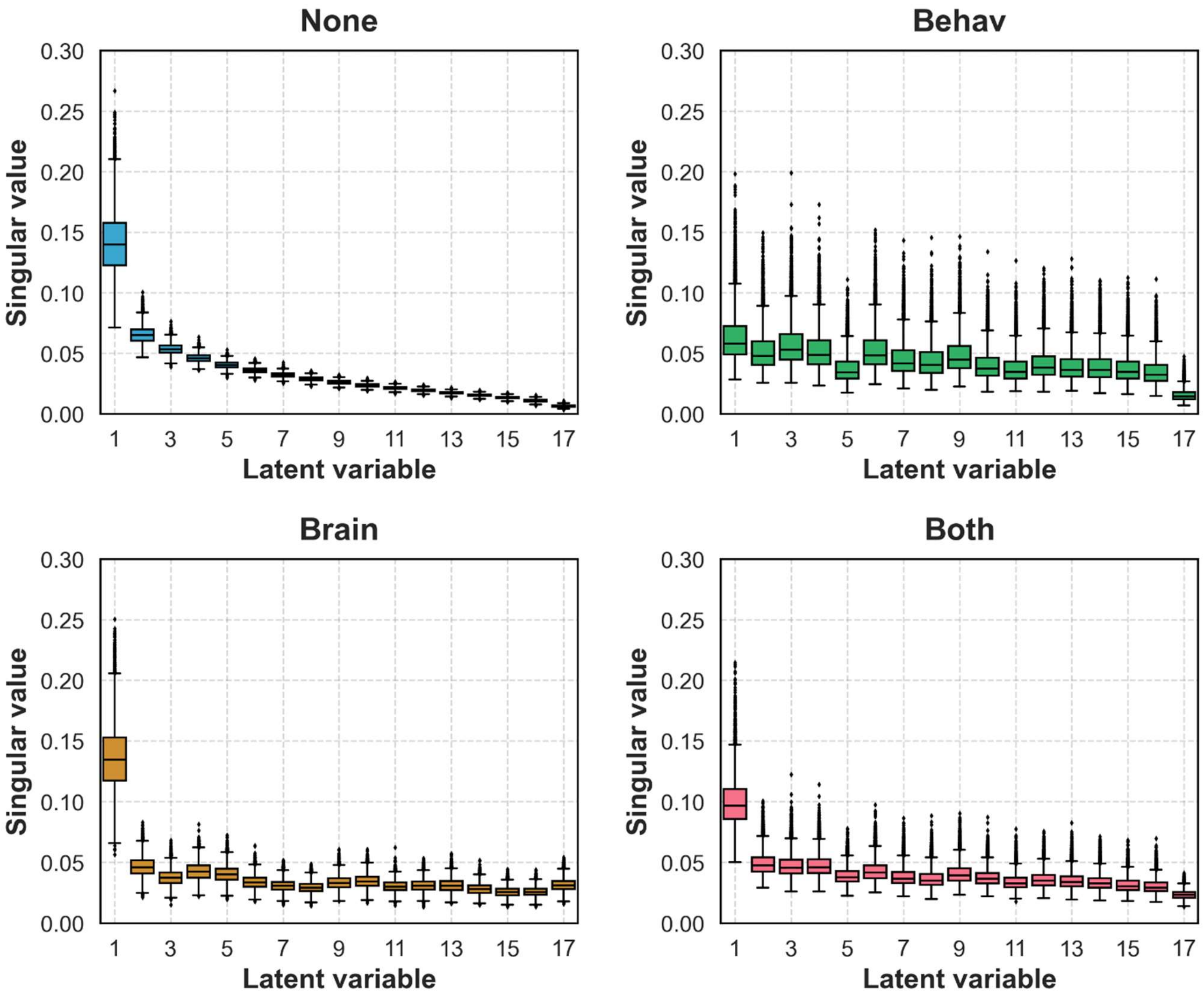


**Figure S12:** null distributions by latent variable for the full UK Biobank sample, plotted separately for each permutation test method. Qualitatively, later null distributions tended to have a higher mean and a greater spread of values after any rotation was applied. In other words, late null distributions appeared to "gain" covariance at the expense of early ones following a rotation. This effect was especially pronounced when the behavioural component was rotated. By extension, when any rotation was applied, early LVs were tested against "weaker" null distributions, and late LVs were tested against "stronger" ones – demonstrating why only the unrotated tests deemed late LVs significant.
